# The plant endoplasmic reticulum is both receptive and responsive to pathogen effectors

**DOI:** 10.1101/2020.06.09.142141

**Authors:** Emily Breeze, Victoria Vale, Hazel McLellan, Laurence Godiard, Murray Grant, Lorenzo Frigerio

**Affiliations:** School of Life Sciences, University of Warwick, Coventry CV4 7AL, UK; Division of Plant Science, University of Dundee (at JHI), Invergowrie, Dundee DD2 5DA, UK; Laboratoire des Interactions Plantes Micro-organismes, CNRS-INRA Toulouse, France

## Abstract

The endoplasmic reticulum (ER) is the entry point to the secretory pathway and, as such, is critical for adaptive responses to biotic stress, when the demand for *de novo* synthesis of immunity-related proteins and signalling components increases significantly. Comprised of a network of interconnected tubules and cisternae, the architecture of the ER is highly pleomorphic and dynamic, rapidly remodelling to meet new cellular requirements. During infection with the hemi-biotrophic phytopathogen, *Pseudomonas syringae* pv. tomato DC3000, the ER in cells immediately adjacent to established bacterial colonies condenses into ‘knot-like’ structures, reminiscent of fenestrated sheets. Based on known temporal dynamics of pathogen effector delivery and initial bacterial multiplication, the timing of these observed morphological changes is rapid and independent of classical elicitor activation of pathogen-triggered immunity. To further investigate a role for ER reconfiguration in suppression of plant immunity we identified a conserved C-terminal tail-anchor domain in a set of pathogen effectors known to localize to the ER and used this protein topology in an *in silico* screen to identify putative ER-localised effectors within the effectorome of the oomycete *Phytophthora infestans*. Subsequent characterization of a subset of 15 candidate tail-anchored *P. infestans* effectors revealed that 11 localised to the ER and/or Golgi. Notably, transient expression of an ER-localised effector from the closely related oomycete, *Plasmopara halstedii*, reconfigured the ER network, revealing intimate association of labelled ER with perinuclear chloroplasts and clusters of chloroplasts, potentially facilitating retrograde signalling during plant defence.

## Introduction

As the gateway to the cell’s secretory pathway, the endoplasmic reticulum (ER) provides the environment for secretory protein production, folding and quality control. The ER is a highly dynamic, interconnected network of tubules and cisternae (sheets) that extends throughout the cytoplasm, associating with the plasma membrane and other organelles, and adjacent cells via plasmodesmata (Hawes et al., 2015). Hence, the ER is central to the maintenance of cellular homeostasis and can facilitate intra- and intercellular communication.

Major transcriptional reprogramming occurs during pathogen infection (Windram et al., 2012; Lewis et al., 2015) and consequently the demand for *de novo* protein and lipid biosynthesis increases significantly. This necessitates rapid but highly regulated ER expansion and/or remodelling, together with an enhanced protein folding capacity to orchestrate successful host defences. However, overloading the ER’s synthesis capacity can result in the accumulation of unfolded and misfolded proteins leading to the unfolded protein response (UPR) and if unmitigated, programmed cell death (PCD) (Srivastava et al., 2018). The UPR is implicated in plant defence via the salicylic acid (SA) dependent signalling pathway which underpins both local and systemic responses to biotrophic pathogens (Wang et al., 2005); and also via SA independent action of the redox-regulated transcriptional cofactor NPR1 (Nonexpressor of Pathogenesis-Related 1) (Wang et al., 2005; Lai et al., 2018). The ER is, therefore, critical to the perception and regulation of adaptive host responses to biotic stress. As a consequence, phytopathogens have evolved ways to target the ER to suppress these immune functions.

Oomycete pathogens such as downy mildews, *Pythium* and *Phytophthora* species infect a wide range of economically important crop and tree species (Kamoun et al., 2015). During infection, oomycetes form specialised structures called haustoria which act as the delivery site for the secretion of both apoplastic and cytoplasmic effectors, and cell-wall degrading enzymes (Wang et al., 2017). In the early stages of pathogen penetration significant cellular reorganisation occurs in the immediate proximity of the haustoria, including the increased association of nuclei and peroxisomes (Boevink et al., 2020); stromule-mediated clustering of chloroplasts (Toufexi et al., 2019) and accumulation of ER and Golgi (O’Connell and Panstruga, 2006; Takemoto et al., 2003). Indeed, the ER itself may be a major source of the extrahaustorial membrane which separates the pathogen from the host cytosol (Kwaaitaal et al., 2017; Bozkurt and Kamoun, 2020).

Genome-wide studies of multiple oomycete species have revealed that they frequently contain large repertoires of the cytoplasmic Arg-X-Leu-Arg (RXLR) class of effectors (Baxter et al., 2010; Haas et al., 2009; Tyler et al., 2006; Jiang et al., 2008; Sharma et al., 2015). These contain an N-terminal signal peptide targeting the protein for secretion, followed by RXLR and EER motifs that are required for subsequent translocation into the host cell (Whisson et al., 2007; Dou et al., 2008). However, the precise route by which cytoplasmic effectors are taken up into the host cell remains unclear.

This arsenal of effectors collectively manipulates multiple host components and signalling pathways to promote virulence (Wang et al., 2019b). Whilst the majority of oomycete RXLRs are targeted to the nucleus (or are dually-targeted to the nucleus and cytoplasm), a subset also localises to the plasma membrane, endomembrane system and chloroplasts (Caillaud et al., 2012; Liu et al., 2018; Pecrix et al., 2019; Wang et al., 2019a).

Experimental validation of ER-localised phytopathogenic effectors and their specific host targets is limited. The *Phytophthora infestans* RXLR effector PITG_03192 has been shown to interact with two potato (*Solanum tuberosum*) NAC transcription factors (TFs) at the ER preventing their translocation to the host nucleus following treatment with *P. infestans* PAMPs (pathogen-associated molecular patterns) (McLellan et al., 2013). These NACs (StNTP1 and 2) are ER-localised via a transmembrane domain (TMD) which, upon signal perception, is proteolytically cleaved allowing translocation of the cytoplasmic domain to the nucleus (Kim et al., 2006; 2010). Arabidopsis contains 14 such annotated NAC with Transmembrane Motif1-like (NTL) TFs of which 12 are validated as being tail-anchored to the ER membrane while NTL5/ANAC060 and NTL11/NAC078 are nuclear-localised (Liang et al., 2015). Besides StNTP1 and 2, other NTLs have also been reported to be targeted by pathogen effectors. These include targeting of NTL9 by the bacterial type III effector HopD1 from *Pseudomonas syringae* (Block et al., 2014) and of LsNAC069 from lettuce (an ortholog of StNTP1) by several effectors from the downy mildew *Bremia lactucae* (Meisrimler et al., 2019). Such interactions demonstrate a theme of diverse pathogen effectors from across the Solanaceae, Brassicaceae and Asteraceae targeting ER tethered NAC TFs as part of conserved host immune suppression strategies.

Aside from preventing the release of ER resident NAC TFs, another pathogen strategy is to deploy effectors that manipulate components of the UPR and thus ER homeostasis. The *Phytophthora sojae* RXLR effector PsAvh262 directly interacts with soybean (*Glycine max*) ER-lumenal binding immunoglobulin proteins (BiPs) - ER quality control chaperones and positive regulators of host susceptibility to selected pathogens including *P. sojae* (Jing et al., 2016). PsAvh262 increases pathogen virulence by stabilizing BiPs and ultimately attenuating ER-stress induced PCD. Similarly, PcAvr3a12 from *P. capsica* directly suppresses the activity of an Arabidopsis ER-localised peptidyl-prolyl cis-trans isomerase (PPIase) involved in protein folding and UPR induction (Fan et al., 2018).

Here, we study how the ER responds to virulent pathogens and their effectors. We first examine ER remodelling during disease development using the well characterised hemi-biotrophic bacterial phytopathogen *Pseudomonas syringae* pv. *tomato* DC3000 (DC3000) infection dynamics. We show that the morphology of the plant ER changes very early in response to a virulent pathogen, and this occurs in a cell-autonomous manner. To highlight the conserved nature of ER targeting by diverse pathogens we test bioinformatic predictions of the intracellular distribution of a group of RXLR effectors from three oomycete species: the economically important *Phytophthora infestans* (causal agent of potato late blight) and sunflower *Plasmopara halstedii* downy mildews, as well as *Hyaloperonospora arabidopsidis* (*Hpa*), a model pathosystem for Peronosporaceae that infects many major crop species (Kamoun et al., 2015). These RXLR effectors all share a similar protein topology: a C-terminal TMD or tail anchor (TA), which in the majority of cases targets them to the ER membrane. We describe a simple and robust *in silico* screening procedure for identifying putative ER- and mitochondrial-targeted proteins within the effectoromes of sequenced pathogen species and validate a subset of these *in planta*. Our results highlight the ER as an important target for pathogen effectors, which drive rapid ER remodelling co-incident with suppression of host defences and initial pathogen multiplication.

## Results

### *Rapid local remodelling of the ER occurs following challenge with the virulent pathogen* Pseudomonas syringae *pv.* tomato *strain DC3000*

The dynamic nature of the ER means that it is constantly remodelling in response to internal and external cues. To assess the temporal and spatial changes in ER morphology in response to a virulent pathogen we utilised the *Arabidopsis thaliana* - *Pseudomonas syringae* pv. *tomato* DC3000 (DC3000) pathosystem, which allows for synchronised infection across a leaf and which we have characterised in detail (Lewis et al., 2015). To this end we used *A. thaliana* expressing the ER luminal marker RFP-HDEL and DC3000 stably expressing eYFP (Rufian et al., 2018), allowing simultaneous visualisation of the ER in the context of the pathogen location. DC3000 effectors are not delivered into the cell until approximately 3 hours post inoculation (hpi) (Lewis et al., 2015), therefore we examined ER morphology in leaf sections by confocal microscopy from 3-10 hpi following DC3000 or mock challenge (10 mM MgCl_2_). No visible changes in ER morphology compared to the Mock treatment were observed up to 6-7 hpi (Figure 1A). By 7 hpi most DC3000 remained planktonic (Figure 1B), although limited immobile bacteria, which deliver type III effectors and pioneer colony establishment, were also becoming established, typically found adpressed between adjacent cells (Bestwick et al., 1995) (Figure 1C-D). At 8 hpi these stationary bacteria begin to multiply (Lewis et al., 2015) as evidenced by initial colony formation (Figure 1E-F). At this timepoint the ER morphology was broadly comparable to that of the mock challenged leaf, however, by 9 hpi substantial remodelling of ER architecture was evident with the ER beginning to condense into tight ‘knot-like’ conformations with the notable loss of resolvable tubular structures (Figures 1G-J). Crucially, the dramatic changes observed in the cortical ER did not occur uniformly across these mesophyll cells, but instead were primarily restricted to those cells immediately adjacent to the establishing bacterial colonies. Distal cells maintained a comparable ER morphology to the mock control (Figure 1J; compare upper cells to the cell at the bottom of the panel). At later timepoints no cortical ER network structure remained.

**Figure 1.**
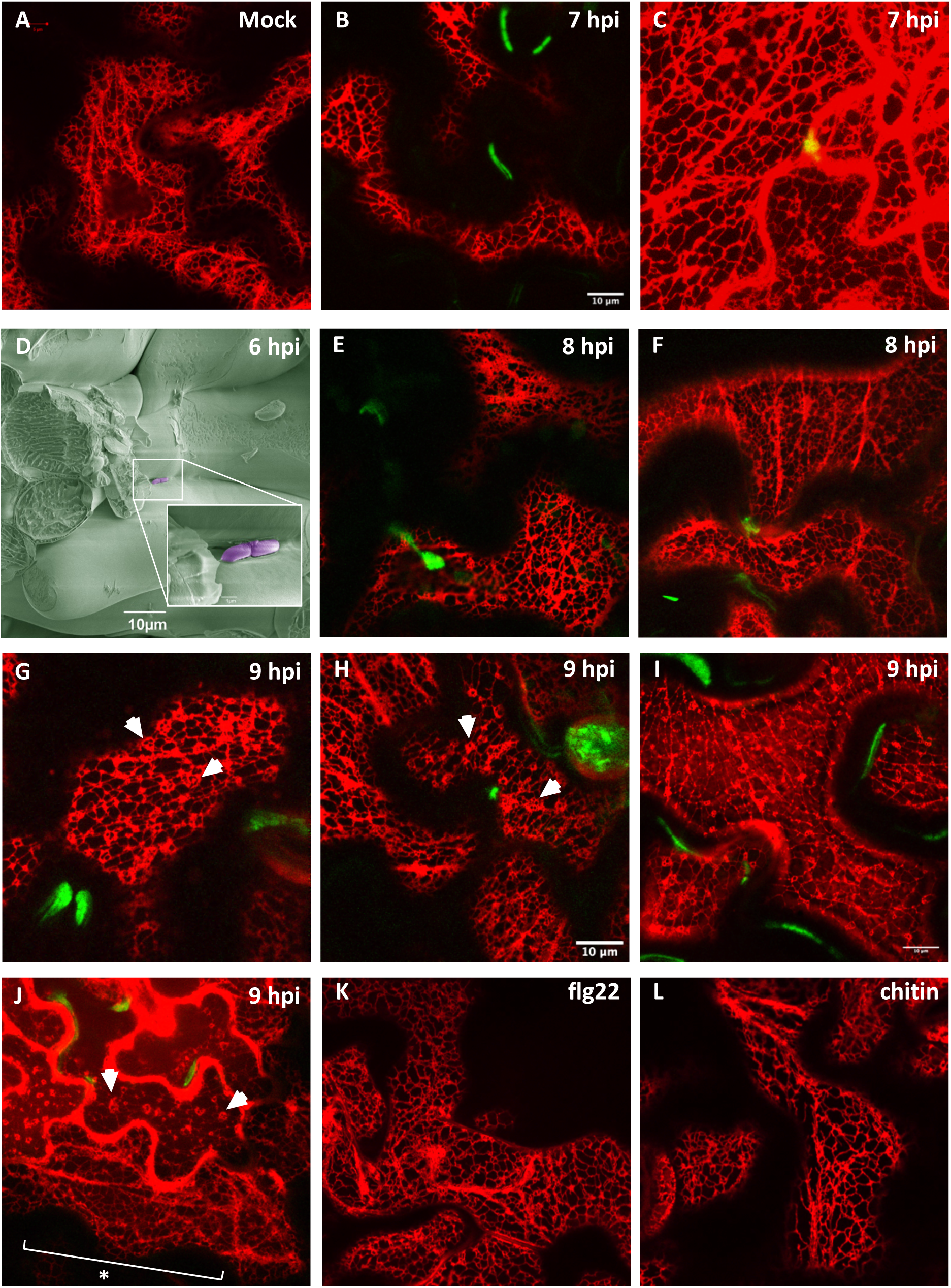
Effector-triggered susceptibility (ETS), but not PAMP-triggered immunity (PTI), initiates ER remodelling and collapse. Representative confocal images of ER labelled with RFP-HDEL (red channel) and eYFP-*Pst* DC3000 (green channel) (excluding D). (A) Mock challenge (6-9 hpi). (B, C) 7 hpi. Bacteria are mainly mobile but small stationary colonies can also be observed between cells. (D) Scanning electron micrograph of *Pst* DC3000 colonies adpressed between two adjacent mesophyll cells (6 hpi). Image taken at 1,300 fold magnification and DC3000 false coloured in purple. Inset shows magnified image (15,000 fold) of boxed area. (E, F) 8 hpi. Discrete bacterial colonies are now evident throughout the tissue. (G-J) 9 hpi. Cells associated with expanding bacterial colonies show disrupted ER morphology with condensed knot-like structures (arrowheads) but this ER phenotype is less visibly affected in distal cells (cell at bottom of image J marked with asterisk). (K-L) Infiltration of known PTI elicitors flg22 (1 µM; K) and chitin (100 µg/ml; L) do not trigger obvious changes in ER morphology cf mock infiltration (Supp Figure 1) 16 hpi.

To test whether whole cell ER remodelling was a function of effector delivery or simply a late PTI (PAMP-triggered immunity) response, we observed leaves infiltrated with the archetypal bacterial or fungal PAMPs. Neither flagellin (1 µM flg22 peptide) nor chitin (100 µg/ml) induced detectable changes in ER architecture up to 16 hpi (Figure 1K-L, Supp Figure 1). Collectively, these data show that virulent DC3000 initiates remodelling of the ER almost co-incident with colony establishment. Moreover, this remodelling is not a generic plant response to infection but rather, it is specifically elicited in a cell-autonomous manner in those cells proximal to bacterial colony establishment. Thus we conclude that ER remodelling is likely invoked by the secretion of bacterial effector molecules into the host, and represents an early and integral part of DC3000’s virulence strategy. Despite the small number of DC3000 effectors, ER targeting has been previously demonstrated for HopD1 by Block et al. (2014); and HopY1 has been predicted to localise to the ER as part of a heteromeric complex comprising the ER localised truncated TIR-plant disease resistance protein, TNL13 and Modifer of SNC1, 6 (MOS6) (Lüdke et al., 2018). However, the observed ER remodelling may also be induced by indirect activities of non-ER targeted effectors.

### The ER is a subcellular target for effectors from both prokaryotic and eukaryotic phytopathogens

Having established that the ER undergoes rapid, gross morphological changes in response to a bacterial pathogen, we next wanted to validate the ER as a target of immune suppression and ascertain whether we could predict the diversity of effectors that target the ER. For this we choose host-pathogen systems with a much more diverse and complex infection strategy: the model pathosystem, *Hyaloperonospora arabidopsidis* (*Hpa*) and two economically important oomycete species, *Phytophthora infestans* and *Plasmopara halstedii* (sunflower downy mildew) – all of which deploy extensive RXLR/RXLR-like effector repertoires. To facilitate this study we first developed a simple bioinformatic screen to predict ER localised effectors.

### C-terminal tail anchor-mediated targeting to the ER membrane is a common strategy employed by oomycete effector proteins

In a large-scale screen Pecrix et al. (2019) characterised a number of RXLR effector proteins expressed by the oomycete *P. halstedii* during infection, of which three, PhRXLR-C13, PhRXLR-C21 and PhRXLR-C22, localised to the ER in *Nicotinia benthamiana* and sunflower transient expression assays. We first confirmed these ER localizations (Figure 2A and Supp Table 1). Despite no significant sequence homology between these three *P. halstedii* effectors, all three are predicted to possess a single transmembrane domain (TMD) positioned towards the C-terminus. Using this observation we examined the predicted topology of a subset of effectors from the closely related oomycete pathogen *Hpa*, which had been previously characterised as localising to the ER when expressed *in planta* (Caillaud et al., 2012). Several of these *Hpa* RXLLs also contained putative TMDs at their C-termini (Figure 2B and Supp Table 1). We thus hypothesised that such tail-anchor (TA) motifs may represent a common ER-targeting mechanism for oomycete effectors, serving to position the effector in the ER membrane with its N-terminus remaining in the cytosol.

**Figure 2.**
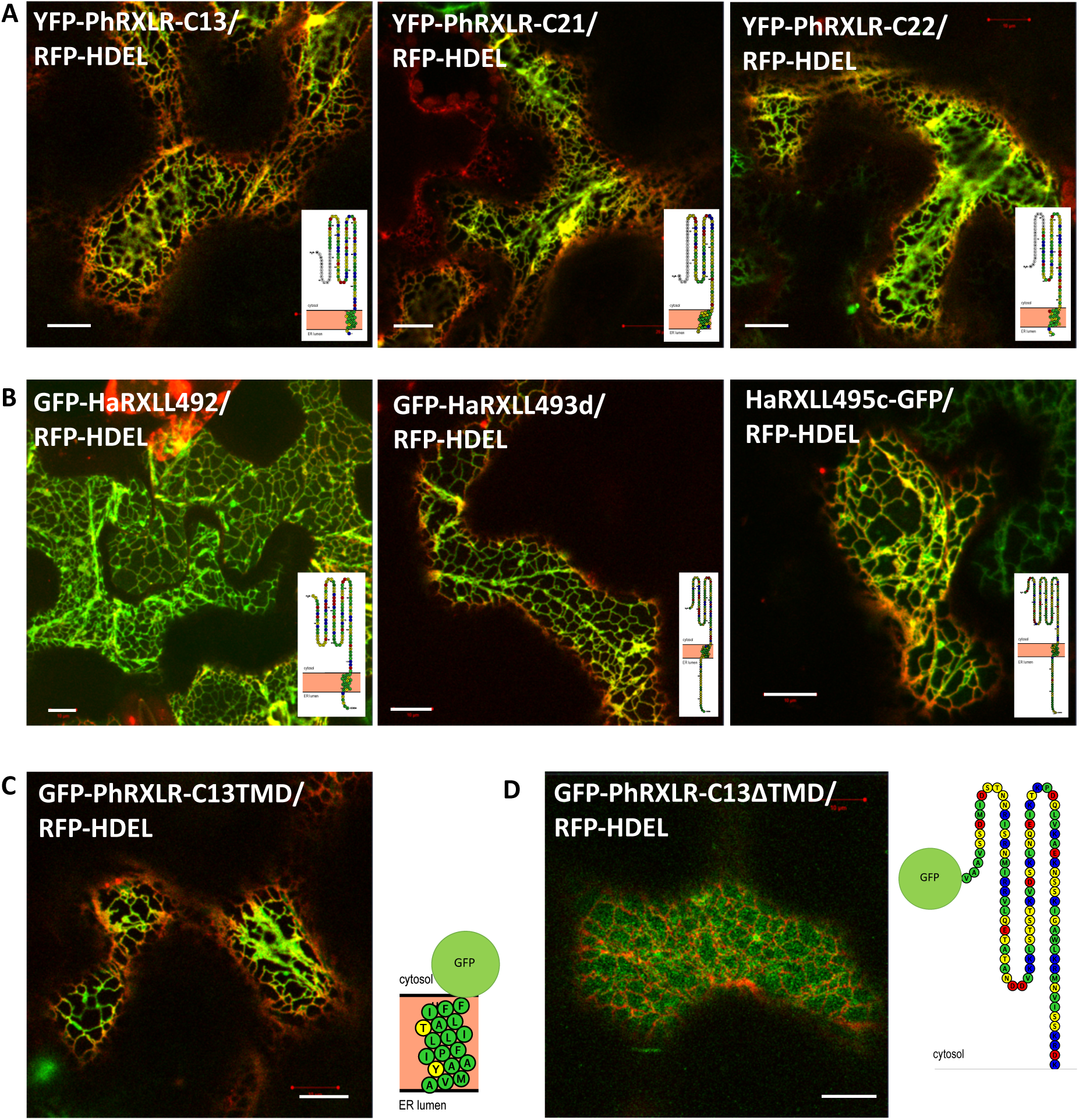
Several ER localised oomycete effectors possess a C-terminal TMD which is sufficient and necessary for ER localisation. Representative confocal images of GFP/YFP-tagged effector proteins (green channel) transiently co-expressed with the ER luminal marker RFP-HDEL (red channel) in *Nicotiana benthamiana* epidermal cells 3 days after infiltration, with TMHMM-predicted protein topology (inset). Scale bar, 10 µm (A) PhRXLR-C13, C21 and C22. (B) HpaRXLL492, 493d and 495a. (C) PhRXLR-C13TMD_108-125._ (D) PhRXLR-C13ΔTMD_108-127_

Using the PhRXLR-C13 effector as an exemplar, we tested whether the TA was required for *in planta* effector localization to the ER. GFP was fused directly to a C-terminal fragment of the PhRXLR-C13 effector consisting of the predicted TA (GFP-PhRXLR-C13TMD_108-125_; Figure 2C). In addition, a truncated version of the effector lacking the TM-spanning region plus the two C-terminal amino acids at the exoplasmic boundary was also generated (GFP-PhRXLR-C13ΔTMD_108-127_; Figure 2D). Whilst GFP-PhRXLR-C13TMD_108-125_ showed comparable ER localization to the full-length fusion protein, GFP-PhRXLR-C13 (Figure 2A), the GFP-PhRXLR-C13ΔTMD_108-127_ lacking the TMD was distributed throughout the cytoplasm Figure 2D). Hence, the presence of a C-terminal TMD is both necessary and sufficient for the ER localization of the PhRXLR-C13 effector.

### Phytophthora infestans has a subset of RXLR effectors with a predicted tail-anchor topology

To test our hypothesis that other oomycete pathogens may also possess a repertoire of ER-targeted effector proteins sharing a similar TA topology, we performed a stringent bioinformatic analysis of the 563 known RXLR effectors from the oomycete *Phytophthora infestans* strain T30-4 (Haas et al., 2009). *P. infestans* is closely related to *P. halstedii* within the Peronosporales order, which also contains *Hpa* (McCarthy and Fitzpatrick, 2017).

We used the membrane topology prediction algorithm TMHMM v2.0 (Krogh et al., 2001) to identify and position any TMDs within the known RXLR effector sequences. TA proteins are inserted post-translationally into their target membrane once the hydrophobic TMD emerges from the ribosome exit tunnel (Hegde and Keenan, 2011). Since this channel is estimated to hold a polypeptide chain of approximately 30 amino acids, the maximal permitted luminal sequence downstream of the predicted TMD was set to 30 residues (Kriechbaumer et al., 2009). Plant ER-localised TM helices are typically between 17-22 residues in length (Brandizzi et al., 2002; Parsons, 2019) and thus an effector was defined as being ‘tail-anchored’ if it possessed a predicted TMD within 50 residues of its C-terminus. These stringent criteria identified 17 putative TA *P. infestans* RXLR effectors, hereafter referred to as Group I effectors (Table 1 and Supp Table 2) and an additional 8 potential candidates (Group II effectors), that fell marginally outside these parameters. The latter comprised 5 effectors with predicted TMDs slightly below the posterior probability cut-off employed by TMHMM and 3 effectors with C-terminal TMDs but beyond the specified final 50 residues. Phylogenetic analysis of (i) the total protein and (ii) TMD sequences of these 25 *P. infestans* TA effectors and the previously characterized *Hpa* and *P. halstedii* ER effectors showed evidence of intra- and inter-species homology, notably in the C-terminal region (Figure 3A). PhRXLR-C13 and the previously characterised PITG_03192 effector (McLellan et al., 2013), for example, have 46% sequence similarity, with HaRxLL492 and PITG_13045 sharing 48% similarity.

**Table 1.**
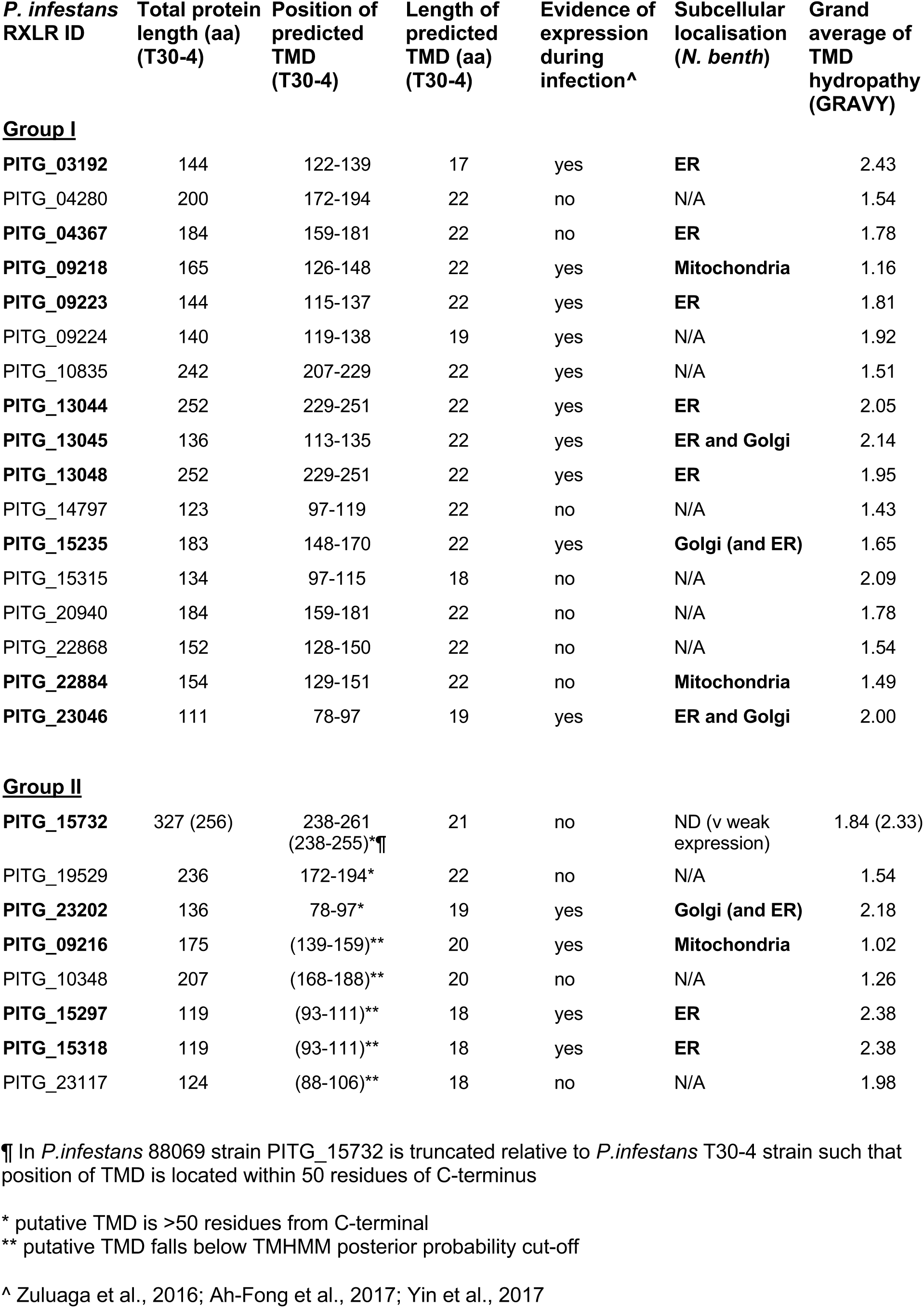
Putative tail-anchored *P. infestans* RXLR effectors. Predicted RXLR effectors from the *P. infestans* T30-4 isolate reference genome (Haas et al., 2009) with putative tail anchors. Subsequent cloning and analyses were performed on effectors derived from *P. infestans* isolate 88069 (Knapova and Gisi, 2002).

**Figure 3.**
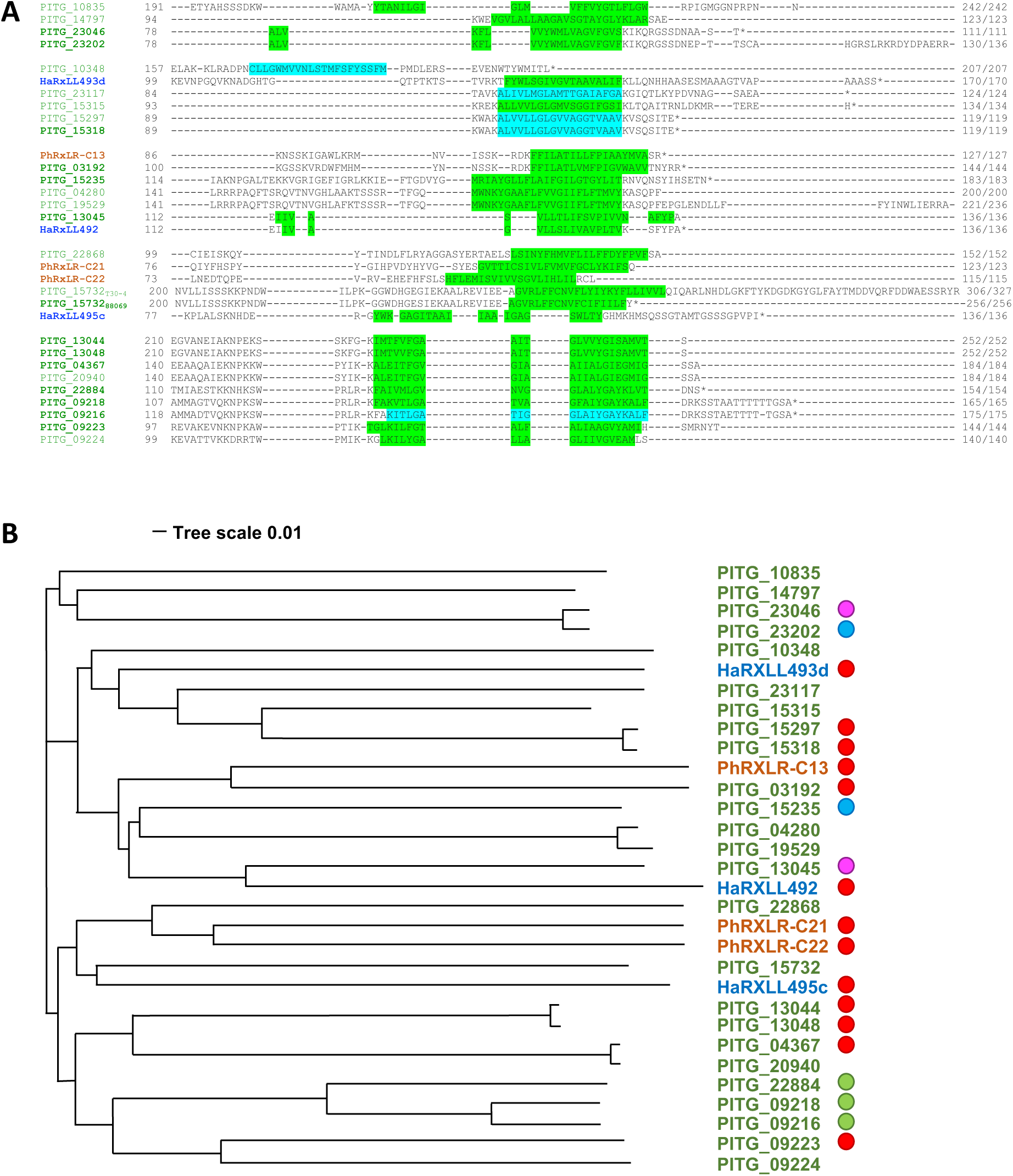
The C-terminal transmembrane domain of tail-anchored effectors are partially conserved between and within oomycete species. (A) Alignment of C-terminal region of *P. infestans* (green), *Hpa* (blue) and *P. halstedii* (orange) effectors within this study. Putative TMDs are highlighted in green (TMHMM prediction) or cyan (TOPCONS prediction). Bold indicates effectors selected for further characterization. (B) Phylogeny of TA effectors from *P. infestans* (green), *Hpa* (blue) and *P. halstedii* (orange) based on whole protein sequences. Filled circles indicate experimentally-determined and/or published (McLellan et al., 2013; Wang et al., 2019a) sub-cellular localization (Red, ER; Blue, Golgi; Pink, ER and Golgi; Green, mitochondria)

We selected a subset of *P. infestans* effectors from both Group I and II for further detailed investigation, ensuring coverage of all the identified phylogenetic clades (Figure 3B). Since the majority of *P. infestans* effectors identified are not experimentally validated we added a criterion for evidence of expression during pathogen infection derived from published RNA-Seq data (Zuluaga et al., 2016; Ah-Fong et al., 2017; Yin et al., 2017). Based upon these conditions, we cloned 10 high confidence (Group I) TA effectors plus 5 Group II effectors (minus the N-terminal signal peptide) (Table 1) from the widely used laboratory isolate 88069 of *P. infestans* (Knapova and Gisi, 2002). As a consequence, some of the cloned sequences exhibited minor amino acid substitutions to the published T30-4 sequences (Supp Table 2), or in the case of PITG_15732 a truncation, resulting in the TMD being positioned within our previously defined TA region. PITG_15732 is a homolog of the well characterized *P. sojae* effector Avr3b, both possessing the nudix hydrolase domain which has been shown to contribute to Avr3b mediated virulence (Dong et al., 2011). While other effectors containing the nudix hydrolase motif are nucleo-cytoplasmic (PITG_06308 and PITG_15679) (Wang et al., 2019a), the presence of the TMD at the C-terminus of PITG_15732 suggested a possible ER address.

### Tail-anchored effectors localize predominantly to the ER and Golgi

To test if the predicted TA effectors localized to the ER *in planta*, we created constitutively expressed N-terminal fluorescent protein-tagged fusions (minus the signal peptide) such that the predicted topology of the chimeric protein had the GFP moiety orientated to the cytosol. Transient expression in *N. benthamiana* epidermal cells and subsequent confocal microscopy 3 days after infiltration allowed subcellular visualisation of the tagged effectors, with the majority exhibiting strong fluorescent protein expression. We could not detect any expression of the PsAvr3b homolog, PITG_15732.

In addition to the six ER-localised tagged *Hpa* and *P. halstedii* effectors, nine of the 15 putative TA *P. infestans* effectors co-localised with the ER luminal marker RFP-HDEL. Two of these effectors (PITG_23046 and PITG_13045) were additionally co-localised with the Golgi marker ST-RFP (Figure 4A and B). A further two effectors, PITG_23202 (highly homologous to PITG_23046) and PITG_15235 were localised mainly to the Golgi (and faintly to the ER). Three of the four remaining GFP-tagged *P. infestans* effectors (PITG_09216, PITG_09218 and PITG_22884) were located to the mitochondria, as evidenced by their co-localisation with the mitochondrial matrix stain, MitoTracker Red™ (Figure 4C) and as previously described by Wang et al. (2019a) for PITG_09218.

**Figure 4.**
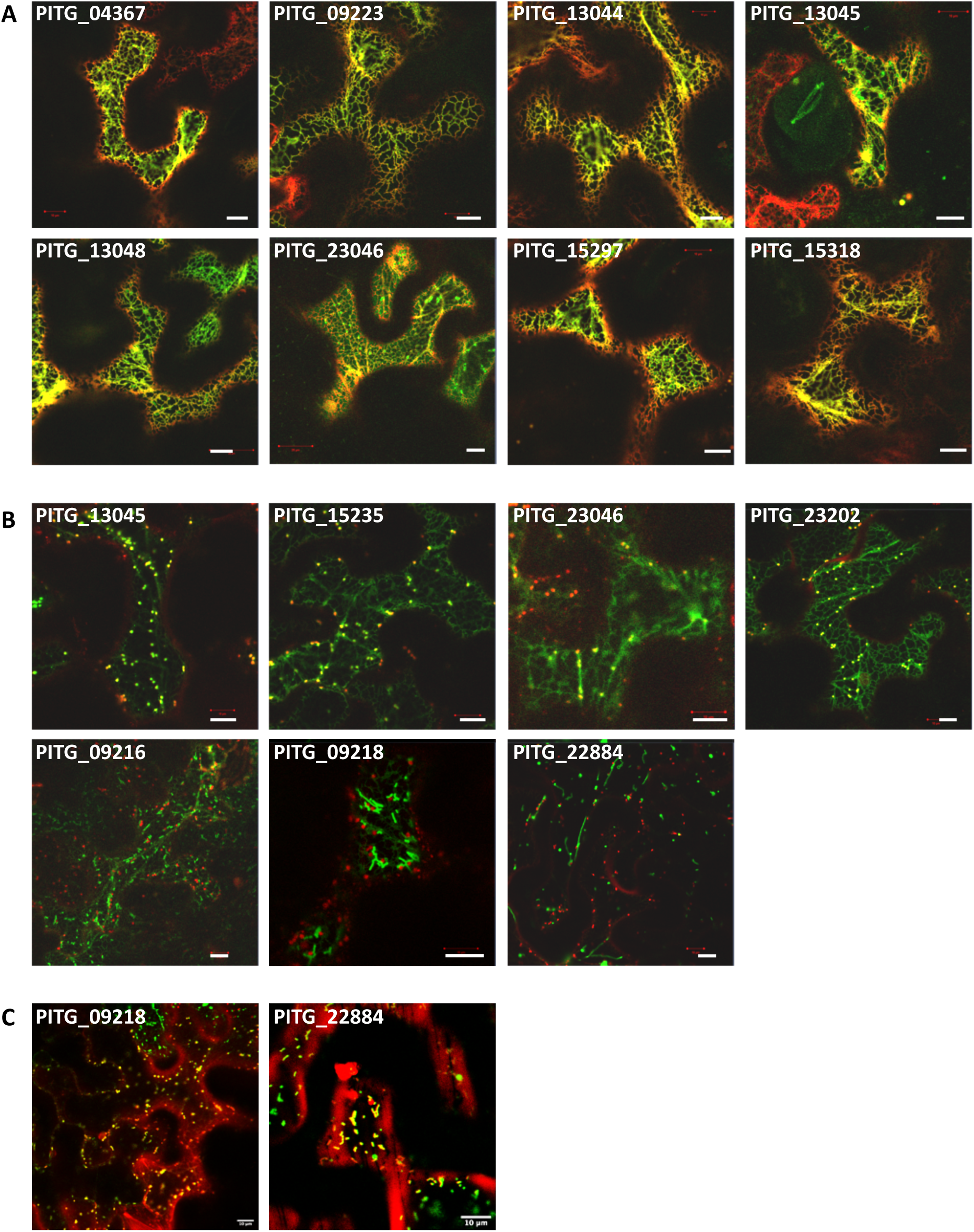
*Phytophthora infestans* tail-anchored RXLR effectors localise to the ER, Golgi and mitochondria. Representative confocal images of 35S:GFP-PITG constructs (green channel) transiently co-expressed in *N. benthamiana* epidermal cells 3 days after infiltration with (A) ER (RFP-HDEL), (B) Golgi (ST-RFP) markers, or (C) post-stained with MitoTracker Red™ (red channel). Scale bar, 10µm.

The precise targeting of TA proteins to their destination membrane depends on multiple physicochemical properties of both the TMD and C-terminal regions. These include the length of the TMD and its hydrophobicity, overall charge of the C-terminal sequence (CTS) and specific motifs therein (Rao et al., 2016; Marty et al., 2014). Here, the length of both the predicted TMD and CTS of the three mitochondrial effectors was comparable to those of the ER-localised effectors (Table 1). Furthermore, although the outer mitochondrial membrane dibasic targeting motif (-R-R/K/H-X^[X≠E]^) (Marty et al., 2014) was present in two of these three mitochondrial localised effectors, it was also present in the ER-localised effectors, PITG_03192 and PITG_23202. The Grand Average of Hydrophobicity (GRAVY) (Kyte and Doolittle, 1982) scores of the *P. infestans* effector TMDs (Table 1) revealed that despite considerable variation in TMD hydrophobicity, the mitochondria-localised proteins had significantly lower values than those of the effectors targeted to the ER and/or Golgi, as previously described (Rao et al., 2016; Kriechbaumer et al., 2009).

### Tail-anchored oomycete effectors converge on membrane-tethered NAC TF targets

Although the specific host protein/s targeted by identified ER-localised effectors have been described in only a handful of cases, several effectors from multiple oomycete and bacterial species converge on the plant NAC with Transmembrane Motif1-like (NTL) family of TFs (Block et al., 2014; McLellan et al., 2013; Meisrimler et al., 2019). To determine if our subset of TA effectors were also capable of interacting with membrane-localised NACs we performed binary yeast two-hybrid (Y2H) assays with 11 of the 14 identified Arabidopsis NTLs (NTL2/ANAC014, NTL5/ANAC060 and NTL9/ANAC116 were not present in our library) (Figure 5).

**Figure 5.**
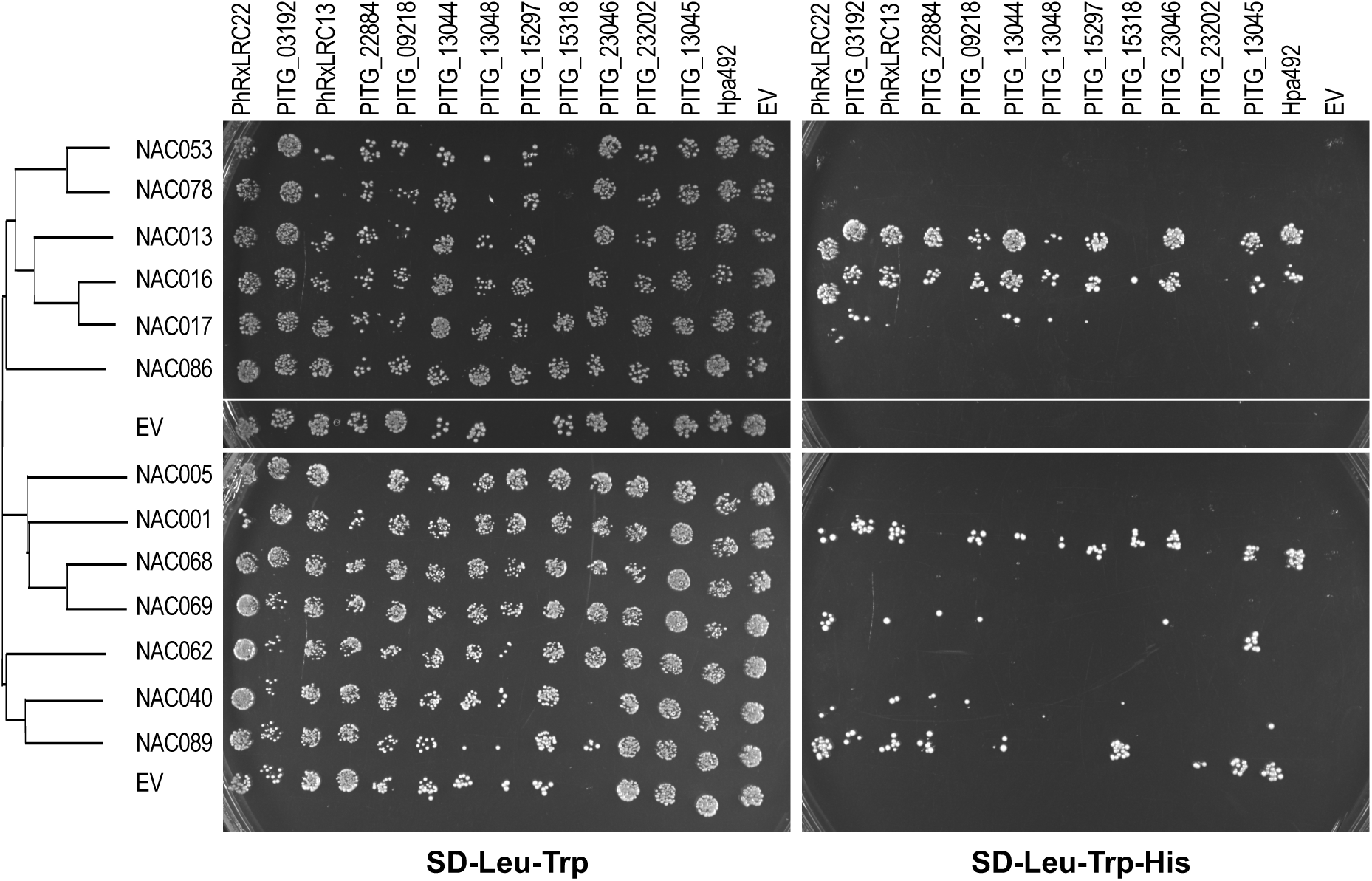
A subset of ER-localised NAC transcription factors interact with several tail-anchored oomycete effectors. Protein-protein interactions between NAC TFs and selected *P. infestans, Hpa* and *P. halstedii* effectors were determined by yeast two-hybrid assays. Positive interaction between bait constructs (effector-GAL4 binding domain fusion) and prey constructs (NAC-GAL4 activation domain fusion) resulting in activation of the HIS3 reporter gene were detected by growth on media lacking histidine (SD-Leu-Trp-His). Growth on SD-Leu-Trp media indicates the presence of both constructs. EV, empty pDEST22 (GAL4 activation domain) or pDEST32 (GAL4 binding domain) vector. For clarity effectors with no detected interactions in replicated assays are not shown.

Several, but not all, of the *P. infestans, Hpa* and *P. halstedii* effectors showed protein-protein interactions with ANAC013 (NTL1), ANAC016 (NTL3) (but not ANAC017 (NTL7)] with which it shares 76% sequence similarity), ANAC001 (NTL10) and ANAC089 (NTL14). This result indicates that multiple ER-directed effectors from phylogenetically diverse pathogens have the capacity to target a specific subset of NTLs, even those of non-adapted hosts. However, other as yet unidentified targets of ER-located effectors are also likely to exist.

### The ER may facilitate perinuclear localisation of chloroplasts during immunity

Possession of a C-terminal TA is likely only one mechanism by which pathogen effectors may be targeted to the ER. HopD1, for example, localises to the ER and targets an ER-localised NTL TF but contains no predicted TMD nor known ER retention motif. We further characterised the localisation of an additional *P. halstedii* RXLR effector, PhRXLR-C20, which was previously described as localising to chloroplasts and stromules (Pecrix et al., 2019). Like HopD1, PhRXLR-C20 contains no predicted TMDs but, in our hands, localised to the ER (Figure 6A). Unexpectedly, however, it also seemed to be tightly associated with chloroplasts (which may explain its previous subcellular assignment) causing them to clump together, notably clustering around the nucleus (Figure 6B, C). We further observed tubular ER with visible polygonal network structure and three-way junctions extending from the ER-wrapped chloroplasts to the nuclear membrane (Figure 6D).

**Figure 6.**
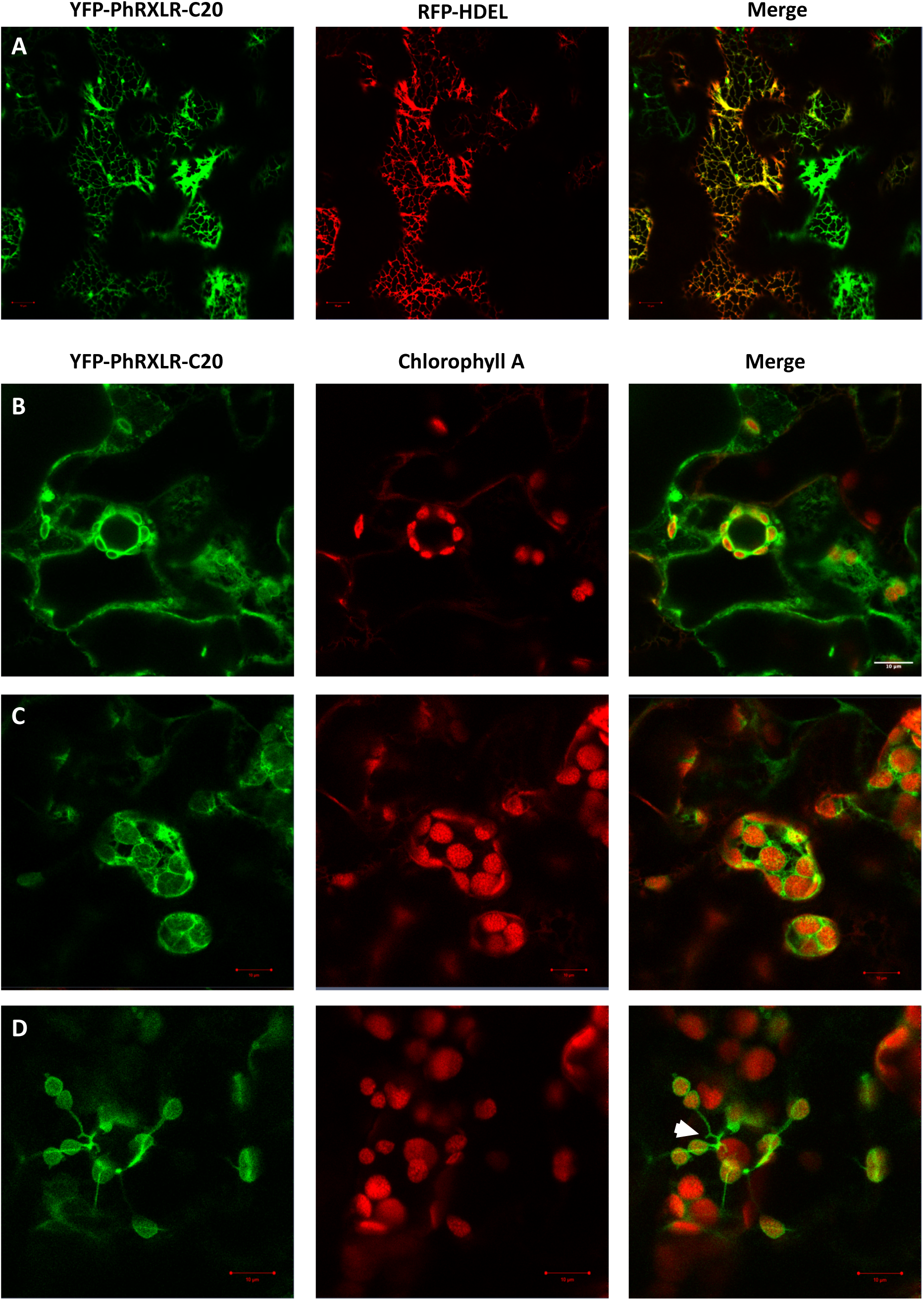
PhRXLR-C20 localises to the ER network in close proximity to chloroplasts. Representative confocal images of YFP-PhRXLR-C20 (green channel) transiently expressed in *N. benthamiana* leaf epidermal cells 3 days after infiltration. (A) YFP-PhRXLR-C20 co-localises with the ER luminal marker RFP-HDEL (red channel). YFP-PhRXLR-C20 labelled ER network appears to wrap around chloroplasts (chlorophyll A-red channel) located both in the cytoplasm (B) and around the nucleus (C). (D) Despite having the appearance of stromules, extensions from the ER-wrapped chloroplasts are likely ER tubules.

## Discussion

In this study we show that gross morphological changes are rapidly manifested in the ER during initial infection with virulent *Pst* DC3000. These changes occur co-incident with initiation of bacterial multiplication (Lewis et al., 2015) and ultimately culminate in the complete collapse of the ER network around 10-12 hpi. We observed that the ER architecture dramatically remodels over a relative short time period of approximately 2 h (7 hpi to 9 hpi) and was evident only in those cells with adjacent bacteria, indicating an early cell autonomous role for DC3000 effectors in ER remodelling. Such morphological changes in ER were not detected in leaves treated with various elicitors of PTI. Local reorganisation of the host ER has also previously been described during infection with various oomycete and fungal pathogens, occurring around both the penetration zone and growing haustoria - the site of effector delivery (Takemoto et al., 2003; Leckie et al., 1995; O’Connell and Panstruga, 2006). Indeed the condensed ‘knot-like’ structures observed here following DC3000 challenge are reminiscent of the perforated, or fenestrated, sheet transitional form of ER visualised around the *P. sojae* infection site (Takemoto et al., 2003). Consequently, this dramatic subcellular event likely represents an early and conserved core pathogen virulence strategy, eventually resulting in the widespread rerouting of the host secretory pathway and the subsequent suppression of plant immunity.

Of the 28 or so effectors in *Pst* DC3000, only HopD1 (Block et al., 2014) has been shown to directly target the ER. However, it was recently reported that HopY1 binds directly to the truncated TIR-NLR, TNL13, possibility facilitating its dissociation from the ER and translocation to the nucleus as a heteromeric complex with MOS6 (Lüdke et al., 2018). Thus the observed changes in ER morphology are likely the result of both direct and indirect responses to manipulation of the ER and/or alteration in the expression of ER-related components by pathogen effectors.

We probed the much larger effectorome of oomycete pathogens using a bioinformatic approach to identify effectors directly targeting the ER. We first identified a subset of ER-localised effector proteins from different oomycete species possessing a highly hydrophobic TMD at their C-termini. An *in silico* screen for the presence of this conserved structural feature within the entire effectorome of *P. infestans* identified several previously uncharacterised effectors targeted to the ER (and Golgi) *in planta*.

Many integral ER membrane proteins possess either a diarginine or dilysine ER retention and retrieval motif (Schutze et al., 1994), whilst soluble ER luminal proteins frequently encode a K/HDEL motif at their C-terminus (Gomord et al., 1997). However, none of the TA effectors characterised in this study contain either of these archetypal ER motifs within their published protein sequence indicating that these effectors have commandeered the TA motif as the primary sorting signal. This was validated using truncated versions of the PhRXLR-C13 effector, to demonstrate that the TMD alone is necessary and sufficient to localise the protein to its target membrane.

TAs are also a known sorting mechanism for proteins resident on the outer envelope of plastids, mitochondria and peroxisomes. Indeed, three of our 15 TA *P. infestans* test effectors were observed to localise to the mitochondria, including PITG_09218 as reported by Wang et al. (2019a). The hydrophobicity of the TMDs clearly discriminates between ER and mitochondrial TA effector TMDs, the latter being weakly hydrophobic (GRAVY<1.5) as previously described in both plant and animal systems (Chio et al., 2017; Rao et al., 2016; Marty et al., 2014; Kriechbaumer et al., 2009). Based on these values we would predict that both PITG_10348 and PITG_14797 localise to mitochondria. Hydrophobicity parameters could thus be incorporated into future iterations of the *in silico* effector screening pipeline to identify likely ER (or mitochondrial) effectors from other pathogen species.

Given the large number of predicted *P. infestans* RXLR effectors (>500), our study identified relatively few ER membrane-localised effectors. The majoirty of tested oomycete effectors localise to the nucleus and/or cytosol with smaller number being targeted to the plasma membrane, chloroplasts, mitochondria and endomembrane system (Caillaud et al., 2012; Pecrix et al., 2019; Wang et al., 2019a; Liu et al., 2018; Khan et al., 2017). However, alternative mechanisms other than the presence of a TA are likely to be employed by effectors targeted to the ER membrane or lumen, such as the aforementioned di-Arg/Lys or H/KDEL motifs. One such candidate is PITG_09585 which encodes the terminal KDEL residues but possesses no predicted TMDs. It is noted that early genome-wide effector discovery pipelines frequently excluded proteins with a predicted TMD (Sperschneider et al., 2015). Consequently, it is feasible that there are additional unannotated ER-localised effectors within the genomes of several well-studied pathogen species.

Effectors need not necessarily embed within the organelle, but rather may associate with the surface or with a resident protein. It is conceivable that effectors targeted to organelles in this manner may function to mark partner binding sites or disrupt membrane contact sites with other organelles, of which the ER forms several (Pérez-Sancho et al., 2016). Identification of such effectors would be of particular interest given the increasing focus in understanding inter-organelle communication in host-microbe interactions and how this is modified by pathogens (Boevink et al., 2020).

Perinuclear chloroplast localisation appears associated with pathogen infection, often accompanied by the extension of stromules towards the nucleus. Chloroplast-nuclear associations have been reported in both avirulent and virulent bacterial challenges, transient expression of viral proteins, exogeneous application of reactive oxygen species and additionally *Agrobacterium tumefaciens* challenge of *N. benthamiana* (Caplan et al., 2015; Erickson et al., 2017; Kumar et al., 2018). Since the ER is an extention of the nuclear envelope, any perinuclear localisation would additionally require negotiation of the ER-nuclear network. Indeed we unexpectedly discovered evidence for a role for the ER in facilitating both chloroplast-chloroplast and perinuclear chloroplast association during transient expression in *N. benthamiana* of the *P. halstedii* effector PhRXLR-C20. In addition to its intimate association with perinuclear chloroplasts, we also observed ER networks coincident with clumping of chloroplasts. Evidence that this is likely an active process is illustrated in Fig. 6D which shows the ER network appearing to ‘draw’ chloroplasts together – somewhat akin to stromules facilitating chloroplast movement to the nucleus (Kumar et al., 2018; Mullineaux et al., 2020).

Pathogenic effectors are under strong selective pressure as part of the perpetual evolutionary arms race with host resistance proteins. However, within the *Phytophthora* genus there is evidence of protein sequence conservation for several effectors, but this is less evident in more distantly related oomycete species. Effector homology is likely indicative of conserved functionality with successful manipulation of the corresponding host target/s being crucial for pathogenicity. Here we identified two pairs of ER localized effectors from different oomycetes, PhRXLR-C13 and PITG_03192; and HpaRXLL492 and PITG_13045, which significant shared sequence homology both outside of and notably within their predicted TMDs.

PITG_03192 localises to the ER in *N. benthamiana* and prevents the relocalisation of two host NAC TFs (NTP1 and 2) from the ER to the nucleus with a corresponding impact on *P. infestans* susceptibility (McLellan et al., 2013). Similarly, Meisrimler et al. (2019) described the interaction of PITG_03192 with a NAC TF from lettuce (*Lactuca sativa*), LsNAC069. LsNAC069 forms a phylogenetic cluster with StNTP2 and with ANAC013, ANAC016 and ANAC017, which were also found to interact with PITG_03192. Here we detected strong interactions with ANAC013 and ANAC016 for both PITG_03192 and the closely related PhRXLR-C13 effector, but a weak, or no interaction, respectively, for these effectors with ANAC017, despite its 76% sequence similarity with ANAC016.

Notably, several of the remaining ER-localised *P. infestans, P. halstedii* and *Hpa* effectors also interacted with ANAC013 and ANAC016, and with ANAC001 and ANAC089, in our Y2H assays. This convergence of multiple effectors on a subset of NAC targets, even in non-adapted pathogens, suggests that these TFs are key players in the host defence response. ANAC013 and ANAC016 are known to be involved in plant tolerance to oxidative stress and drought conditions, respectively (De Clercq et al., 2013; Sakuraba et al., 2015) with ANAC016 also implicated in the regulation of leaf senescence (Kim et al., 2013).

During ER stress ANAC089 relocates from the ER to the nucleus in a bZIP28- and bZIP60-dependent manner, promoting the transcriptional upregulation of genes associated with the UPR and PCD (Yang et al., 2014). Several transcriptional regulators within the UPR pathway (many of which are ER-membrane associated in non-activated conditions) serve as points of convergence with known environmental stress signalling pathways including bZIP28, bZIP60, NF-YA4 and NF-YC2, and NPR1 (Liu and Howell, 2010; Moreno et al., 2012; Lai et al., 2018). The ER quality control system may therefore act as an early sensor and signal transducer of environmental stress conditions, enabling the ER secretory machinery to be primed to meet the increased demand for stress-related proteins (Pastor-Cantizano et al., 2020). Hence, the UPR is critical for adaptive immune responses with the direct or indirect manipulation of various components of the UPR pathway by effectors likely representing a common virulence strategy employed by pathogens.

In high-throughput and stringent Y2H screens, ANAC089 formed protein-protein interactions with HopD1 (previously demonstrated to be ER localised (Block et al., 2014)]) and with VAP27-1 (Mukhtar et al., 2011). VAP27-1 also interacted with two of the Hpa effectors (HpaRXLL492 and HpaRXLL495) described in this study. VAP27-1, together with NET3C mediates the formation of contact sites between the ER and the plasma membrane (Wang et al., 2016; 2014). As discussed above, ER contact sites are an attractive target for pathogen manipulation in order to derail intracellular communications during infection, and may account for the ‘knotted’ ER appearance induced by DC3000 infection. However, attempts to confirm the Y2H interaction between VAP27-1 and the two *Hpa* effectors by both co-immunoprecipitation and FRET-FLIM analysis were unsuccessful (data not shown).

For several of the ER-localised effectors we did not detect interactions with members of the NTL TF family in our Y2H assays, indicative of other potential ER protein targets. Y2H assays are context-free and thus the detected effector-NAC interactions need to be confirmed by alternative methods and their biological relevance investigated *in planta*. This lack of biological context is well illustrated by the observed interactions of the mitochondrial PITG_09218 and PITG_22884 effectors with the ER-localised ANAC013, ANAC016, ANAC001 and/or ANAC089 TFs.

In summary, we have demonstrated that the ER undergoes a rapid and radical reconfiguration during *Pst* DC3000 infection, coincident with the initiation of bacterial colony establishment and the known temporal dynamics of effector delivery into the host cell. Based on this observation we used a bioinformatic approach to identify a number of effectors from multiple oomycete species that are targeted to the host ER by virtue of possession of a C-terminal tail anchor. Whilst the presence of a signal peptide targets the effectors for secretion via the conventional secretory pathway, it remains unclear how such membrane-associated proteins are subsequently trafficked to their host target, which presumably requires shielding or masking of the hydrophobic TMD during translocation.

It is becoming increasingly clear that effectors possess cellular addresses other than the nucleus and cell wall, with an increasing focus on suppression of chloroplast immunity (de Torres-Zabala et al., 2015). We propose that the ER, as the major site of *de novo* lipid and protein biosynthesis, is also a prime target for manipulation by multiple pathogens orchestrated through the secretion of a suite of diverse effectors specifically targeted to this organelle.

## Materials and Methods

### Plant materials and growth conditions

*Arabidopsis thaliana* stably expressing the ER luminal marker RFP-HDEL, were grown for 4-5 weeks in a compost mix (Levingston F2) in a controlled environment growth chamber programmed for 10 h day (21°C; 120 μmol m^−2^ s^−1^) and 14 h night (21°C) regime with 60 % relative humidity. *Nicotiana benthamiana* were grown for 5-7 weeks under a 16 h day (21°C) and 8 h night (18°C) regime.

### Constructs and plant transformation

Candidate tail-anchored *P. infestans* effectors were cloned without their signal peptides as predicted by SignalP (Armenteros et al., 2019). Sequences were amplified from *P. infestans* isolate 88069 (Knapova and Gisi, 2002) genomic DNA using gene-specific primers flanked with a portion of the Gateway *att*B recombination sites (all primer sequences are given in Supp Table 3). A second round of PCR was performed with full length *att*B primers with the resulting *att*B-PCR product purified and used to generate an entry clone in pDONRZeo. N-terminal sGFP fusions of the effectors were created by performing an LR recombination reaction with the Gateway binary destination vector pGWB606 (Nakamura et al., 2014). This cloning strategy was also used to generate a truncated version of PhRXLR-C13 lacking the predicted TMD (GFP-PhRXLR-C13ΔTMD_108-127_), and a GFP-fusion of the predicted PhRXLR-C13 TMD alone (GFP-PhRXLR-C13TMD_108-125_).

All effector constructs and organellar marker plasmids (RFP-HDEL [ER] or ST-RFP [Golgi]) were transformed via heat-shock into *Agrobacterium tumefaciens* strain GV3101 and were transiently expressed into *N. benthamiana* leaf epidermal cells at an OD_600_ of 0.2, as previously described (Sparkes et al., 2006). Leaf cells were imaged 3 days after infiltration.

### Bacterial growth, maintenance and inoculation

*P. syringae* pv. *tomato* strain DC3000 expressing eYFP (Rufian et al., 2018) was grown on solidified Kings B medium containing 50 µg ml^-1^ rifampicin and 25 µg ml^-1^ kanamycin. For inoculation, cells from an overnight culture grown at 28°C were harvested by centrifugation at 2800 *g* for 7 min, washed and resuspended in 10 mM MgCl_2_. Cell density was adjusted to an OD_600_ of 0.15. Mature, upper rosette Arabidopsis leaves were infiltrated with the bacterial suspension or 10 mM MgCl_2_ (Mock) using a 1 ml needleless syringe on their abaxial surface. For PTI elicitor treatment, chitin (100 µg/ml) or flg22 peptide (1 µM) were infiltrated into an independent leaf in an identical manner. Leaf cells were imaged 3-10 h (DC3000) or 16 h (chitin and flg22) post infiltration.

### Microscopy and imaging

#### Confocal microscopy

Freshly excised leaf samples were mounted in water and imaged on a Zeiss LSM 880 confocal microscope with a Plan-Apochromat 100× (DC3000) or 63x (effectors)/ 1.40 oil DIC M27 objective. GFP was excited at 488 nm and detected in the 498-563 nm range; mRFP was excited at 561 nm and detected in the 602-654 nm range; chlorophyll A was excited at 561 nm and detected in the 605-661 nm range. Mitochondria were stained with 100 nM MitoTracker™ Red (Invitrogen), washed in water and imaged after 10-60 min.

#### Electron microscopy

Infected leaf samples were removed by razor blade and mounted upright onto a cryo-sledge coated using a 1:1 mix of OCT compound / colloidal graphite and rapidly frozen in liquid nitrogen slush (Alto 2100 cryo system, Gatan, Ametek, Leicester, UK). The frozen samples were then transferred under vacuum into the cryo-pre-chamber. To reveal the cellular interior of the leaves a movable blade within the cryo-chamber was used to produce a freeze-fracture. Water was sublimated for 3 min at -95°C followed by sputter-coating with gold/palladium (80/20) within the cryo-pre-chamber. Samples were imaged at -135°C using a JEOL 6390LV scanning electron microscope operated at 2 kV and micrographs subsequently false-coloured in Adobe Photoshop CS6.

### Yeast-2-hybrid assays

GAL4 DNA-binding domain fusions were generated for all *P. infestans, P.halstedii* and *Hpa* effectors in this study by recombination with pDEST32 (Invitrogen) and subsequent transformation of the bait construct into the haploid Y8930 (MATα) yeast strain. A Y2H prey library of Arabidopsis NTL proteins fused to the GAL4 activation domain (pDEST22; Invitrogen) was similarly created and transformed into the opposite yeast mating strain, Y8800 (MATa). Yeast-2-hybrid assays were performed as described in (Harvey et al., 2020). Empty pDEST22 and pDEST32 vectors (EV) transformed into Y8800 and Y8930 yeast strains, respectively, were used as negative controls.

### *In silico* analysis of RXLR effectors

Predictions of the membrane topology of RXLR effectors, notably the position and length of the TMD, were performed using both the TMHMM v2.0 (Transmembrane prediction using Hidden Markov Model) (Krogh et al., 2001) and TOPCONS (Tsirigos et al., 2015) algorithms. All annotated *P. infestans* RXLR effector sequences were screened to identify putative TA proteins based on the presence of a single TMD 17-22 residues in length located at the C-terminal with a maximum of 30 residues permitted after the predicted TMD. In fact over half of the predicted TA effectors identified in this study had less than 10 residues post-TMD.

### Phylogenetic analysis

Protein sequences of selected effectors were aligned using Clustal Omega (Sievers et al., 2011) and a phylogenetic tree generated using iTOL (Interactive Tree of Life) (Letunic and Bork, 2019).

### Accession numbers

ANAC001, NTL10, AT1G01010; ANAC005, AT1G02250; ANAC013, NTL1, AT1G32870; ANAC014, NTL2, AT1G33060; ANAC016, NTL3, AT1G34180; ANAC017, NTL7, AT1G34190; ANAC040, NTL8, AT2G27300; ANAC053, NTL4, AT3G10500; ANAC060, NTL5, AT3G44290; ANAC062, NTL6, AT3G49530; ANAC068, NTL12, AT4G01540; ANAC069, NTL13, AT4G01550; ANAC078, NTL11, AT5G04410; ANAC086, AT5G17260; ANAC089, NTL14, AT5G22290; ANAC116, NTL9, AT4G34480.

## Supporting information

Support Tables 1-3

## Supplemental Material

**Supplemental Figure 1.**
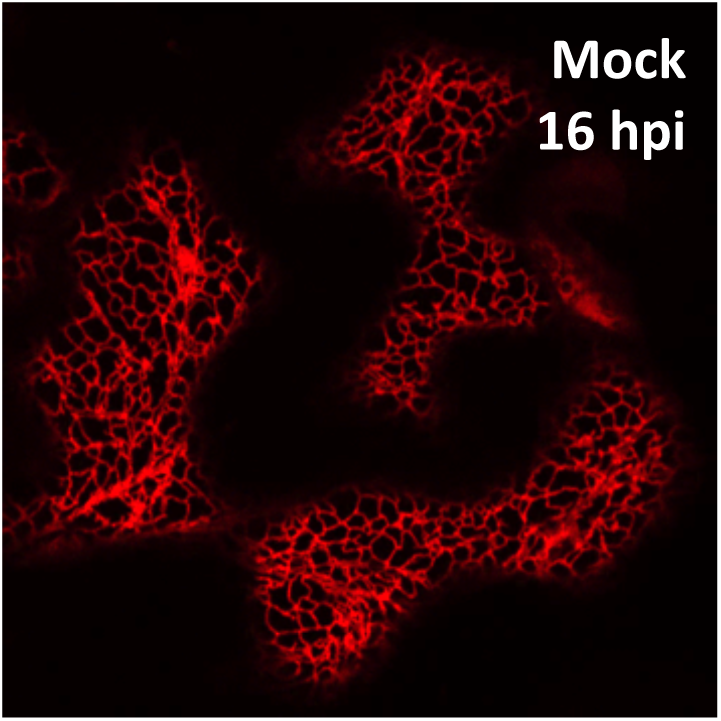
Mock infiltration control (16 hpi). ER morphology is comparable to that of leaves infiltrated with known PTI elicitors flg22 (1 µM; Figure 1K) and chitin (100 µg/ml; Figure 1L).

**Supplemental Table 1.** Selected tail-anchored effectors from *Plasmopara halstedii* and *Hyaloperonospora arabidopsidis* characterised in this study

**Supplemental Table 2.** Putative tail-anchored effectors from *Phytophthora infestans* identified in this study.

**Supplemental Table 3.** Primers used in this study

## Acknowledgements

We thank Petra Boevink (JHI at University of Dundee) for critical evaluation of the manuscript; Yann Pecrix (CIRAD) for preparation of *P. halstedii* effector constructs; Laurence Tomlinson (TSL) for the kind donation of the Hpa RXLL effector constructs, and Christian Hacker (Exeter) for assistance with SEM. This work was supported by internal University of Warwick funding to LF. MG acknowledges support from BBSRC/UKRI grant BB/P002560/1.

*In memory of Chris Hawes who ignited a love of cell biology in so many.*

## Author Contributions

EB, MG and LF conceived and designed the study. EB and VV performed all experiments. EB performed the data analysis. LG cloned and undertook initial characterisation of the *P. halstedii* effectors. EB, LF, HM and MG prepared the manuscript.

